# The length of lipoteichoic acid polymers controls *Staphylococcus aureus* cell size and envelope integrity

**DOI:** 10.1101/2020.03.23.004671

**Authors:** Anthony R. Hesser, Leigh M. Matano, Christopher R. Vickery, B. McKay Wood, Ace George Santiago, Heidi G. Morris, Truc Do, Richard Losick, Suzanne Walker

## Abstract

The opportunistic pathogen *Staphylococcus aureus* is protected by a cell envelope that is crucial for viability. In addition to peptidoglycan, lipoteichoic acid (LTA) is an especially important component of the *S. aureus* cell envelope. LTA is an anionic polymer anchored to a glycolipid in the outer leaflet of the cell membrane. It was known that deleting the gene for UgtP, the enzyme that makes this glycolipid anchor, causes cell growth and division defects. In *Bacillus subtilis*, growth abnormalities from the loss of *ugtP* have been attributed to the absence of the encoded protein, not to loss of its enzymatic activity. Here, we show that growth defects in *S. aureus ugtP* deletion mutants are due to the long, abnormal LTA polymer that is produced when the glycolipid anchor is missing from the outer leaflet of the membrane. Dysregulated cell growth leads to defective cell division, and these phenotypes are corrected by mutations in the LTA polymerase, *ltaS*, that reduce polymer length. We also show that *S. aureus* mutants with long LTA are sensitized to cell wall hydrolases, beta-lactam antibiotics, and compounds that target other cell envelope pathways. We conclude that control of LTA polymer length is important for *S. aureus* physiology and promotes survival under stressful conditions, including antibiotic stress.

**IMPORTANCE:** Methicillin-resistant *Staphylococcus aureus* (MRSA) is a common cause of community- and hospital-acquired infections and is responsible for a large fraction of deaths caused by antibiotic-resistant bacteria. *S. aureus* is surrounded by a complex cell envelope that protects it from antimicrobial compounds and other stresses. Here we show that controlling the length of an essential cell envelope polymer, lipoteichoic acid, is critical for controlling *S. aureus* cell size and cell envelope integrity. We also show that genes involved in LTA length regulation are required for resistance to beta-lactam antibiotics in MRSA. The proteins encoded by these genes may be targets for combination therapy with an appropriate beta-lactam.

## INTRODUCTION

The bacterial cell envelope is a barrier that protects bacteria from unpredictable and often hostile environments. In Gram-positive bacteria, such as *Staphylococcus aureus*, the cell envelope comprises the cell membrane and a thick peptidoglycan (PG) layer that is decorated with a variety of proteins and polymers important for viability and virulence. Among these polymers are teichoic acids, which are negatively charged and divided into two classes based on their subcellular localization. One class, wall teichoic acids (WTA), are covalently linked to PG; the other class, lipoteichoic acids (LTA), are associated with the cell membrane through a glycolipid anchor (1). WTA and LTA play partially redundant roles in cell envelope integrity and cannot be deleted simultaneously (2-4).

In *S. aureus*, both WTA and LTA have been implicated in the control of cell morphology and division (3, 5, 6), virulence (7-14), osmoregulation (15-18), antimicrobial resistance (6, 19-22), and spatiotemporal regulation of cell wall enzymes (23-27). However, LTA is more important than WTA for cell viability. *S. aureus* can grow under standard laboratory conditions without WTA (28), but cells lacking lipoteichoic acid synthase (LtaS), the enzyme that assembles LTA on the cell surface, are not viable and rapidly acquire suppressor mutations (3, 5, 16, 29, 30).

The usual glycolipid anchor and the starting unit for LTA is diglucosyl-diacylglycerol (Glc_2_DAG), which is synthesized from UDP-glucose and diacylglycerol (DAG) by the glycosyltransferase UgtP (also called YpfP) (Fig. 1A-B) (31, 32). Glc_2_DAG is exported to the cell surface by LtaA (10). The lipoteichoic acid polymerase is LtaS, a polytopic membrane protein with an extracellular domain that contains the active site (4, 33). LtaS transfers phosphoglycerol units derived from phosphatidylglycerol (Ptd-Gro) to the Glc_2_DAG starter unit, producing DAG as a byproduct (5, 34, 35). DAG is recycled to Ptd-Gro by a salvage pathway (36). LTA is heavily decorated with D-alanyl residues, a modification of teichoic acids that is important in autolysin regulation and has also been implicated in *S. aureus* virulence (8, 12, 13, 19). Notably, deleting the glycosyltransferase gene *ugtP* or the genes encoding the enzymes that produce UDP-glucose, its substrate, does not result in the loss of LTA, although such mutations result in morphological and fitness defects (10, 11, 32). Instead, LtaS uses Ptd-Gro rather than Glc_2_DAG as the lipid starter unit for LTA assembly.

**FIG 1.**
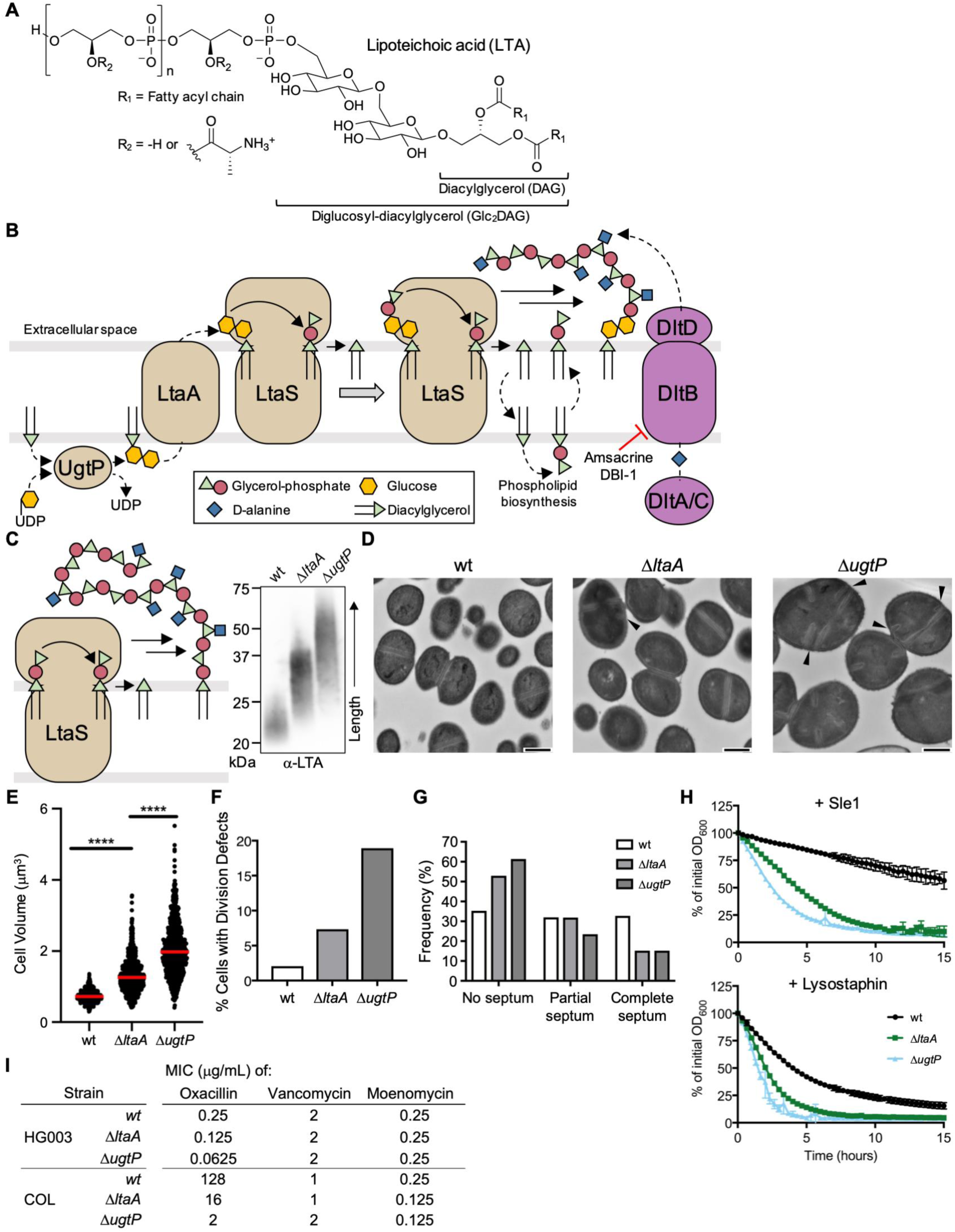
*S. aureus* mutants unable to synthesize LTA on Glc_2_DAG produce long LTA polymers anchored on Ptd-Gro, have cell size and division defects, and are susceptible PG hydrolases and beta-lactam antibiotics. **A)** LTA is a polymer of 30-50 phosphoglycerol repeats linked to a Glc_2_DAG lipid anchor. Repeat units are heavily modified with D-alanyl esters. **B)** UgtP uses DAG and UDP-glucose to synthesize Glc_2_DAG, which is then exported by LtaA to the cell surface. LtaS polymerizes phosphoglycerol units derived from Ptd-Gro on the lipid anchor. Each elongation cycle generates DAG. The *dlt* pathway adds D-alanyl esters to LTA. **C)** In the absence of Glc_2_DAG, LtaS uses Ptd-Gro as an alternative lipid anchor, resulting in abnormally long LTA (see anti-LTA Western blot of exponential phase RN4220 lysates). **D)** TEM of RN4220 strains. Arrowheads indicate septal defects. Scale bar = 500 nm. **E**) RN4220 cell volumes were calculated from cells lacking visible septa. The median is shown with a red bar. **** indicates p < 0.0001. **F)** RN4220 cells were classified based on the cell cycle stage and presence of defects (see FIG S1C for classification scheme). Cells containing misplaced and/or multiple septa (classes C and E) were totalled. **G)** Cell cycle stage frequency plot of classified cells without visible defects. **H)** RN4220 strains were suspended in phosphate-buffered saline and treated with lytic enzymes. The decrease in OD_600_ was tracked over time. For all plots, the average and standard deviation from 3 technical replicates are shown. **I)** MIC table of oxacillin, vancomycin, and moenomycin against select *S. aureus* strains. HG003 is an MSSA strain and COL is an MRSA strain.

In *Bacillus subtilis*, deleting *ugtP* or the genes for the enzymes that produce UDP-glucose also results in morphological defects (37-40). In one study, *ugtP* mutant cells were shown to be shorter than wild type cells, and it was proposed that UgtP is a nutrient sensor that negatively regulates cytokinesis in a manner that depends on UDP-glucose levels (38). In this model, under nutrient-rich conditions, high levels of UDP-glucose localize UgtP to the cytokinetic ring and inhibit FtsZ polymerization or constriction, which delays cell division to provide cells time to grow to a larger size (38, 41). Δ*ugtP* cells are therefore small because UgtP is not present to slow cell division. Other studies in *B. subtilis* are not consistent with a role for UgtP in nutrient sensing because mutant cells were found to be enlarged in its absence or to have shape rather than size alterations (39, 40).

UgtP evidently does not act to increase cell size in *S. aureus* because Δ*ugtP* mutant cells are larger, rather than smaller, than those of the wild type (32). Like other *S. aureus* cells that grow too large (6, 42-45), the mutant cells have cell division defects such as multiple and misplaced septa (32). While these defects may result directly from the loss of the *ugtP* gene product, the Δ*ugtP* deletion is pleiotropic and causes multiple effects, including the absence of the disaccharide anchor Glc_2_DAG and an abnormal lengthening of LTA polymers (Fig. 1C, Fig. S1A) (10). Whether increased cell size and dysregulated cell division directly result from the loss of the *ugtP* gene product, from the loss of intracellular Glc_2_DAG, or from the abnormally long LTA polymers has been unclear.

Mutants lacking *ltaA*, which encodes the flippase that exports Glc_2_DAG to the cell surface, provide a means to distinguish among these possibilities. Δ*ltaA* mutants express UgtP and synthesize intracellular Glc_2_DAG, but are unable to export it efficiently (10). The LTA polymers produced by Δ*ltaA* mutants are longer than those of the wild type but shorter than those of Δ*ugtP* mutant cells (Fig. 1C), being a heterogeneous mixture in which some polymers are assembled on Ptd-Gro and some on Glc_2_DAG (which is exported to the cell surface by an alternative, unknown mechanism) (10). We hypothesized that if the defects observed in Δ*ugtP* cells were caused by the abnormally long LTA polymers, we should observe alterations in cell size and division in Δ*ltaA* mutant cells, although perhaps less pronounced due to the intermediate polymer length caused by the Δ*ltaA* mutation.

Here we show that the production of long, abnormal LTA is sufficient to alter cell size and lead to cell division defects. We also report that LTA pathway mutants with these morphological defects are highly susceptible to beta-lactam antibiotics and PG hydrolases and are dependent on other cell envelope pathways that are dispensable in wild type strains. We used an inhibitor of one of these pathways to select for suppressor mutations in Δ*ugtP* strains and found that most of the suppressor mutations were located in the LTA polymerase, *ltaS*, and caused a reduction in LTA polymer length. Polymer abundance was frequently decreased as well, in some cases to almost undetectable levels. The *ltaS* suppressor mutations partially reversed the cell size and division abnormalities caused by the Δ*ugtP* mutation. Taken together, these studies indicate LTA length and abundance plays a crucial role in controlling cell size and cell envelope integrity in *S. aureus*.

## RESULTS

### Δ*ugtP* and Δ*ltaA* mutants are larger than wild type cells and have cell division defects

A previous study in *S. aureus* reported that Δ*ugtP* cells are larger than wild type cells and have other morphological defects (32), but Δ*ltaA* mutant cells have not been examined in detail. We compared the morphology of otherwise isogenic wild type, Δ*ltaA*, and Δ*ugtP* strains by transmission electron microscopy (TEM) and quantified cell size using brightfield and epifluorescence microscopy after staining with a membrane dye. Compared with wild type, Δ*ltaA* cells appeared larger by TEM and Δ*ugtP* cells were clearly larger (Fig. 1D). Quantification of cell size confirmed these observations: cell volume increased from 0.72 ± 0.14 μm^3^ for wild type to 1.26 ± 0.38 μm^3^ for Δ*ltaA* and 1.98 ± 0.59 μm^3^ for Δ*ugtP* (Fig. 1E, Fig. S1B). Δ*ltaA* and Δ*ugtP* cells also displayed cell division defects, including multiple and misplaced septa (Fig. S1C-D). Only 2.1% (n = 536) of wild type cells had these defects compared with 7.3% (n = 518) for Δ*ltaA* cells and 18.9% (n = 55) for Δ*ugtP* cells (Fig. 1F). Furthermore, Δ*ltaA* and Δ*ugtP* cells were more commonly observed without partial or complete septa, indicating that they spend a longer time growing prior to initiating septal synthesis (Fig. 1G, Fig. S1E). These shared phenotypes of Δ*ltaA* and Δ*ugtP* mutants indicate that control over cell growth and division is adversely affected when LTA is abnormally long and assembled on a Ptd-Gro, rather than on a Glc_2_DAG, membrane anchor. An important challenge for the future will be to elucidate the mechanism by which LTA polymer length influences cell size and division.

### Δ*ugtP* and Δ*ltaA* mutants are more susceptible to lytic enzymes than wild type

Given the established connection between teichoic acids and enzymes that act on the cell wall, we used Δ*ltaA* and Δ*ugtP* mutants to probe whether production of abnormal LTA causes susceptibility to lytic enzymes that degrade peptidoglycan. Sle1 is an amidase native to *S. aureus* that hydrolyzes the amide bond between N-acetylmuramic acid and the stem peptide of PG (46). Lysostaphin is an endopeptidase produced in other *Staphylococci* spp. that hydrolyzes the peptide cross bridges between PG strands (47, 48). We found that Δ*ltaA* and Δ*ugtP* mutants were substantially more susceptible to these enzymes than were wild type cells (Fig. 1H, Fig. S1F), indicating that cells that make long, abnormal LTA have alterations in their cell envelope that cause increased susceptibility to enzymes that hydrolyze PG.

### Δ*ugtP* and Δ*ltaA* MRSA mutants are sensitized to beta-lactam antibiotics

Previous work has shown that when WTA is removed, methicillin-resistant *S. aureus* (MRSA) becomes susceptible to some beta-lactam antibiotics, including oxacillin (6). WTA-null cells also have cell size and division defects that resemble those that we observed for Δ*ltaA* and Δ*ugtP* (6). These observations prompted us to test whether deleting *ltaA* or *ugtP* in a MRSA background also affected beta-lactam susceptibility. Although the minimum inhibitory concentration (MIC) for oxacillin was only modestly decreased when *ltaA* or *ugtP* was deleted in a methicillin-sensitive background, it decreased by 8-fold and 64-fold for Δ*ltaA* and Δ*ugtP* mutants, respectively, in MRSA (Fig. 1I, Fig. S1G). By contrast, there was little to no change in the MICs of antibiotics that primarily affect peptidoglycan polymerization (moenomycin and vancomycin) rather than crosslinking. The susceptibility of Δ*ltaA* and Δ*ugtP* to beta-lactams provides further evidence that these mutants have cell wall alterations.

### Δ*ugtP* and Δ*ltaA* mutants lyse when teichoic acid D-alanylation is blocked

We found in previous work that Δ*ugtP* cells become reliant on a tailoring modification of LTA called D-alanylation in which D-alanyl esters are attached to the C2 hydroxyl groups of the phosphoglycerol repeats of LTA (49). This modification, although dispensable in wild type strains, is important in resistance to cationic antimicrobial peptides and aminoglycosides (15, 19), and has been linked to regulation of autolysin function (50). We previously identified two structurally unrelated inhibitors, DBI-1 and amsacrine (amsa), that inhibit DltB, an essential component of the D-alanylation pathway (Fig. 1B, Fig. 2A) (49, 51). Using spot titre assays, we found here that Δ*ltaA* strains, like Δ*ugtP* strains, are susceptible to both DltB inhibitors (Fig. 2B, Fig. S2A). This susceptibility provides additional evidence that the production of abnormal, long LTA causes defects that result in conditional essentiality of other cell envelope modifications.

**FIG 2.**
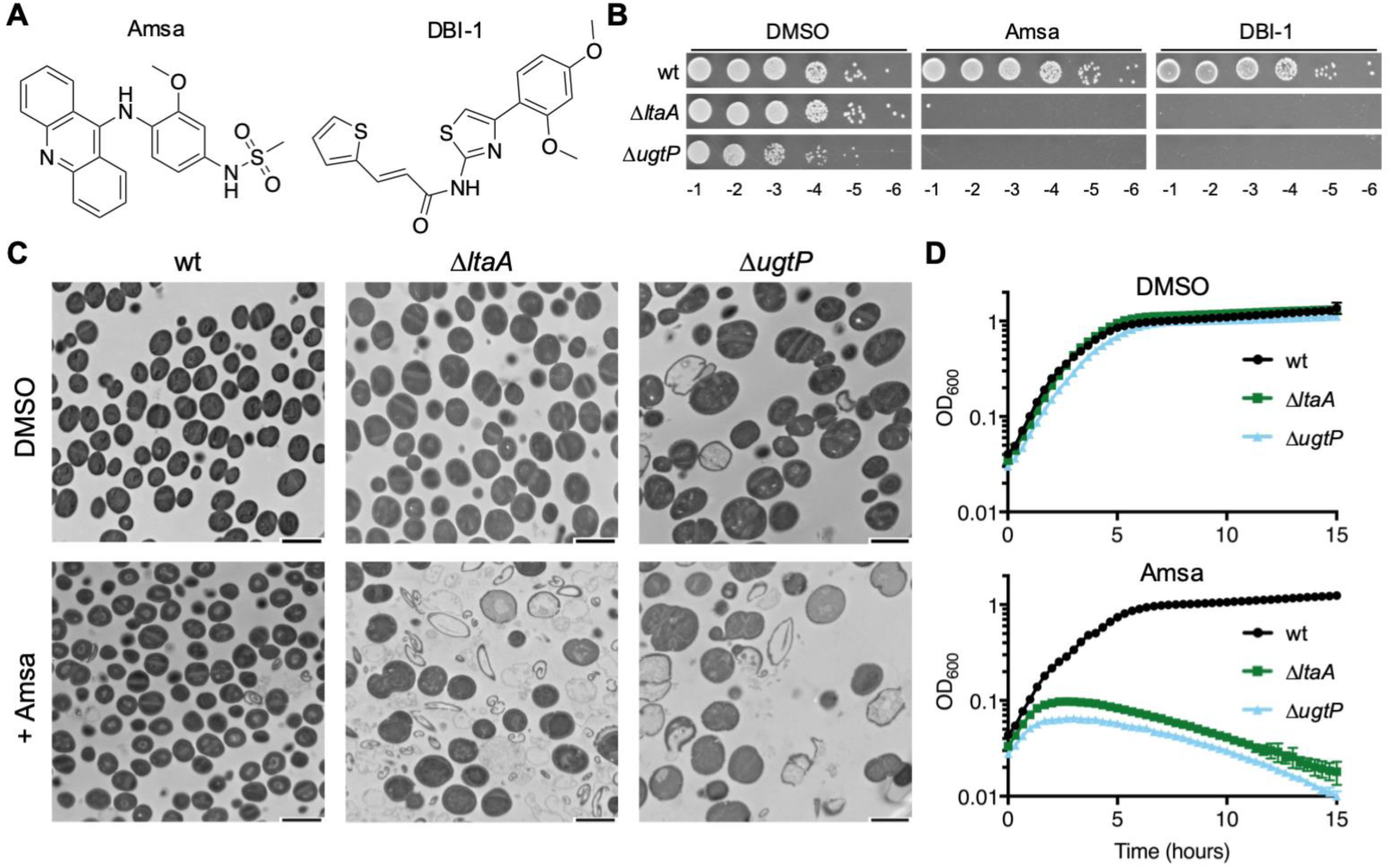
Mutants with long, abnormal LTA undergo lysis when treated with D-alanylation inhibitors. **A)** Structures of the two DltB inhibitors used in this study. **B)** 10-fold serial dilutions of RN4220 strains on 10 μg/mL DltB inhibitor or DMSO vehicle control. **C)** TEM of RN4220 strains treated with DMSO or with 10 μg/mL amsa. Scale bar = 2 μm. **D)** Growth curves of RN4220 strains treated with DMSO or 10 μg/mL amsa. The average and standard deviation from 3 technical replicates are shown.

Terminal phenotypes can provide insight into processes that lead to cell death. We therefore used TEM to compare the phenotypes of wild type, Δ*ltaA*, and Δ*ugtP* cells treated with the DltB inhibitor amsa. Fields of untreated and treated wild type cells appeared similar by TEM, but there were large numbers of cell ghosts and fragments in images of the treated Δ*ltaA* and Δ*ugtP* mutants (Fig. 2C, Fig. S2B). Moreover, when we monitored biomass by optical density after treatment with inhibitor, we observed that the mutants stopped growing and then the optical density dropped (Fig. 2C, Fig. S2C-D). The TEM images of the untreated Δ*ugtP* mutants also showed qualitatively more lysis than wild type cells. Taken together, our findings show that Δ*ltaA* and Δ*ugtP* cells have a greater propensity to lyse than wild type cells, and they undergo catastrophic lysis when D-alanylation is inhibited. We infer that D-alanylation negatively regulates the activity of cell wall hydrolases.

### Reducing LTA length and abundance suppresses lethality of D-alanylation inhibitors

Because suppressor analysis can provide clues to the underlying basis of a defect, we selected for suppressor mutations of amsa lethality in a Δ*ugtP* strain. Whole genome sequencing of one suppressor showed a mutation in *ltaS* that resulted in a leucine substitution at F93, and we found that LTA produced by this mutant was shorter and less abundant than in the parent strain (see below). We raised additional amsa-resistant mutants from multiple independent cultures in a few distinct *ugtP* null strains, all of which could be complemented (Fig. S3A). Targeted sequencing of *ltaS* showed that mutations in this gene accounted for 75% of the 64 resistant colonies tested. Notably, only three of the mutations resulted in premature stop codons. In all other mutants, the reading frame of *ltaS* was maintained, suggesting that at least partial function of the encoded protein was retained.

To assess whether the activity of LtaS was affected by the *ltaS* mutations, we selected 14 strains that contained mutations spanning different regions of the encoded enzyme (Fig. 3A, Fig. S3B) and evaluated the abundance and length of their LTA. LtaS contains five predicted transmembrane (TM) helices and a C-terminal extracellular domain that contains the active site (4, 33). Three mutants contained 18 base-pair insertions that resulted in the insertion of six amino acids in the extracellular loop between TM1 and TM2. The other eleven mutants contained single amino acid substitutions in another extracellular loop, in the “helical linker” region, or in the extracellular domain. For the mutants for which LTA was detectable by Western blot, we found that it was shorter than that in the *ugtP* parent strain and was typically much less abundant (Fig. 3B). For some mutants, LTA was undetectable. Targeted sequencing did not identify suppressor mutations known to restore viability to *ltaS* knockouts (16, 29, 30). Moreover, whole genome sequencing of a subset of strains did not identify any shared mutations in other genes (Fig. S3C). Taken together, these results suggested that suppression of the lethality caused by loss of D-alanylation in Δ*ugtP* was a direct result of the reduced LTA length and abundance.

**FIG 3.**
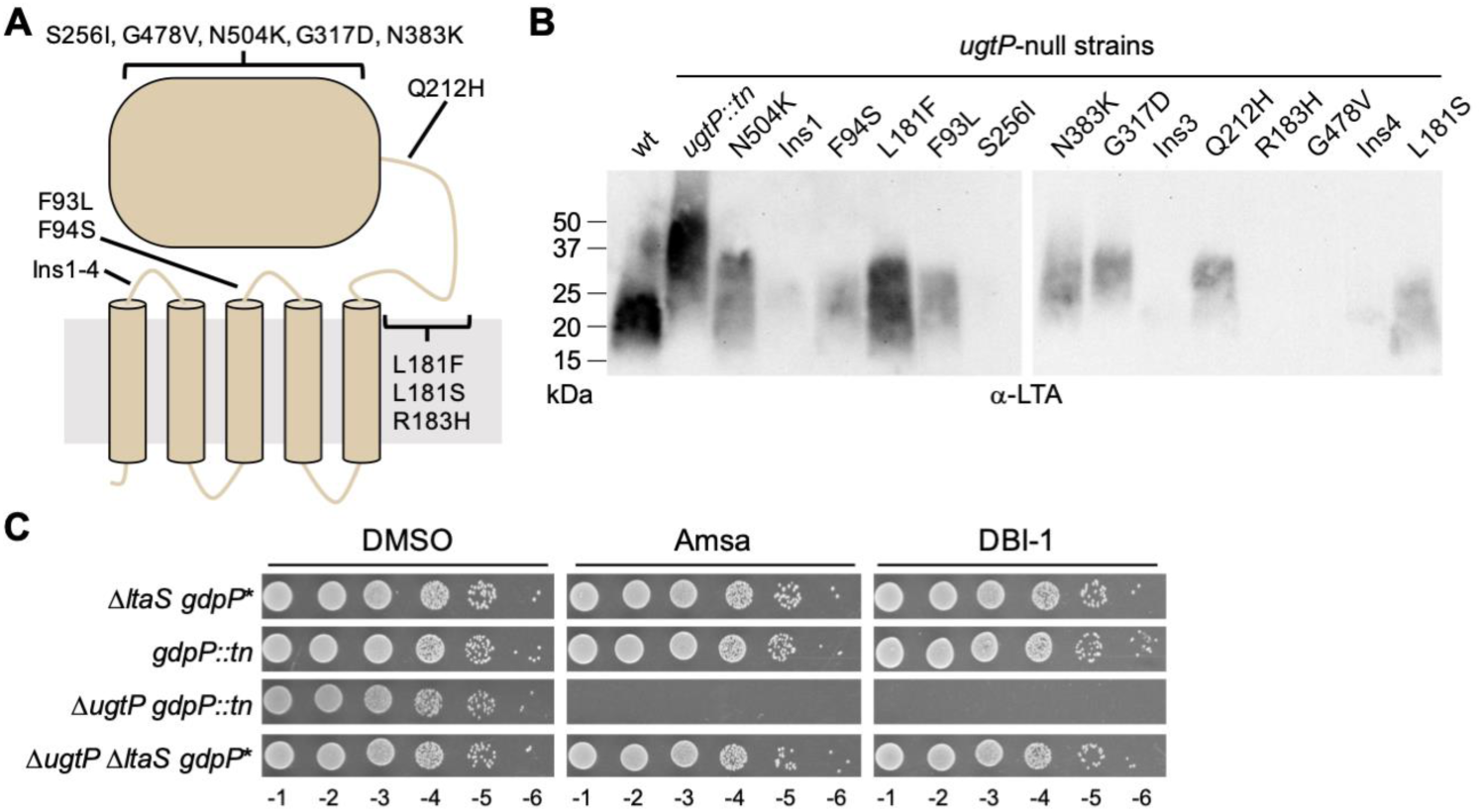
The lethality of amsa to Δ*ugtP* is suppressed by mutations in LtaS that decrease LTA length and/or abundance. **A)** Schematic of LtaS depicting the location of the identified mutations. **B)** Anti-LTA Western blot of exponential phase lysates from isolated *ltaS* mutants in *ugtP* null backgrounds. While LTA abundance cannot be precisely compared between LTA of different lengths with the anti-LTA antibody, several mutations appear to drastically decrease LTA abundance. The wild type and *ugtP::tn* strains are in the SEJ1 background, while the strain background of each isolated mutant is given in FIG S3B. **C)** 10-fold serial dilutions of strains on plates containing 10 μg/mL DltB inhibitor or DMSO. The Δ*ltaS* strain contains a *gdpP*\* hypomorphic suppressor mutation. *gdpP::tn* is a transposon disruption of *gdpP*. Δ*ltaS* strains are in the SEJ1 background, while the *gdpP::tn* strains are in the RN4220 background.

Consistent with this conclusion, we found that a Δ*ugtP* Δ*ltaS* double deletion strain was not susceptible to amsa because it does not produce LTA (Fig. 3C). This strain contains a hypomorphic mutation in *gdpP* (*gdpP*\*). GdpP is a cyclic-di-AMP phosphodiesterase and this mutation results in increased cellular levels of cyclic-di-AMP, a second messenger implicated in osmoregulation (52). The accumulation of cyclic-di-AMP allows growth in the absence of *ltaS* (16). We therefore confirmed that a Δ*ugtP gdpP*::Tn double mutant remained sensitive to amsa and produced LTA resembling that of a Δ*ugtP* single mutant (Fig. 3C, Fig. S4A-B). Therefore, amsa resistance in the Δ*ugtP* Δ*ltaS* double mutant was not due to *gdpP*\*, but rather to the absence of LTA.

To verify that amsa resistance in the suppressor strains was indeed caused by the *ltaS* mutations, we constructed strains expressing inducible copies of three *ltaS* mutant alleles (Ins3, F93L, and L181S) or the wild type allele. These mutations are found in extracellular loops of the transmembrane domain (Ins3, F93L) or in the linker region between the transmembrane domain and the extracellular domain (L181S). We then deleted the chromosomal copy of *ltaS* and introduced marked deletions of *ltaA* or *ugtP* into each strain (Fig. 4A). The strains required inducer for growth, indicating that the mutant *ltaS* alleles, like the wild type allele, performed an essential function (Fig. 4A). Strains expressing the *ltaS* mutants grew on amsa and DBI-1 regardless of whether *ltaA* or *ugtP* were present; however, strains expressing *ltaS*^*wt*^ in a Δ*ltaA* or Δ*ugtP* background did not grow (Fig. 4A, Fig. S5A-B). Growth curves showed that amsa-treated cells expressing *ltaS*^*wt*^ in the Δ*ugtP* background underwent growth arrest followed by lysis, while cells expressing the *ltaS* mutants continued to grow (Fig. 4B). For two of the mutants, *ltaS*^*F93L*^ and *ltaS*^*L181S*^, growth rates in amsa were comparable to the untreated controls, but the *ltaS*^*Ins3*^ mutant was growth-impaired. This mutant also reached only 20% of the optical density of the other mutants in overnight cultures. Despite differences in fitness, the reconstructed mutants confirmed that the selected *ltaS* mutations are necessary and sufficient to confer resistance to D-alanylation inhibitors in a Δ*ugtP* background.

**FIG 4.**
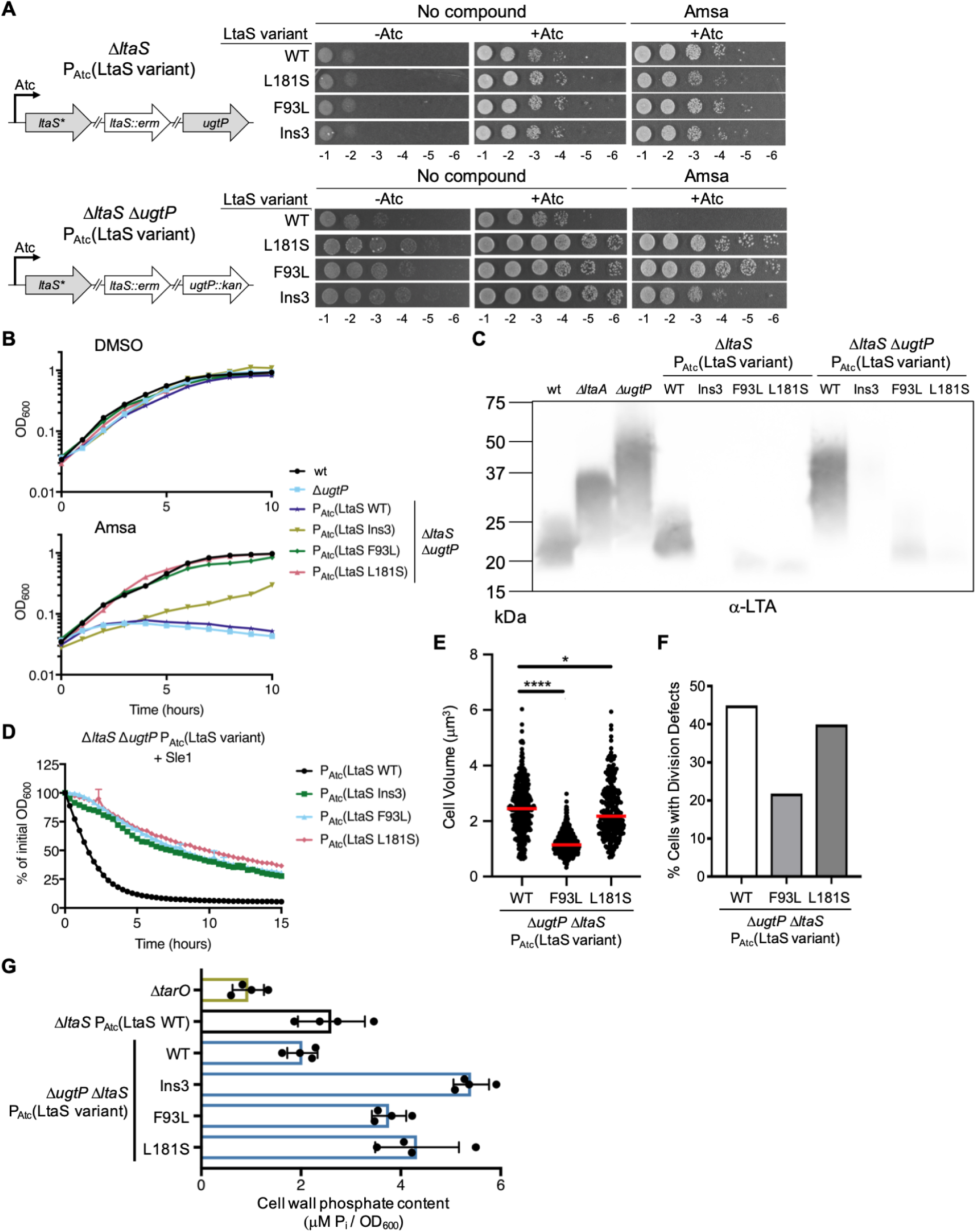
Mutations in *ltaS* that reduce LTA length and/or abundance correct defects of Δ*ugtP* and increase WTA abundance. **A)** RN4220 with an anhydrotetracyline (Atc)-inducible copy of select *ltaS* mutants was transduced with an erythromycin (erm)-marked *ltaS* deletion and, optionally, a kanamycin (kan)-marked *ugtP* deletion. Rebuilt suppressor strains were spotted in a 10-fold dilution series on plates containing 10 μg/mL amsa or DMSO. Note that some residual growth is observed on plates without inducer due to growth of cells in the presence of inducer prior to plating. **B)** Growth curves of RN4220 Δ*ugtP* strains with ectopically expressed *ltaS* variants as the only source of LtaS. Strains were treated with 10 μg/mL amsa or DMSO. A representative growth curve from 3 replicates is shown for each strain. **C)** Anti-LTA Western blot of exponential phase lysates from rebuilt *ltaS* mutants. **D)** RN4220 Δ*ugtP* strains expressing *ltaS* alleles were suspended in phosphate-buffered saline and the decrease in OD_600_ was tracked upon addition of Sle1. This phenotype is also observed with lysostaphin and when expressing *ltaS* mutant alleles in the wild type background (FIG S7). The average and standard deviation from 3 technical replicates are shown. **E)** Cell volumes of reconstructed RN4220 Δ*ugtP* strains expressing *ltaS* alleles were calculated from cells lacking visible septa. Expression of *ltaS*^*F93L*^ (1.15 ± 0.31 μm^3^, median ± median absolute deviation) or *ltaS*^*L181S*^ (2.17 ± 0.78 μm^3^) in the Δ*ugtP* background reduces size compared to expression of *ltaS*^*wt*^ (2.45 ± 0.73 μm^3^). The median is shown with a red bar. * indicates p < 0.05 and **** indicates p < 0.0001. **F)** RN4220 Δ*ugtP* strains expressing *ltaS* alleles were classified based on cell cycle stage and presence of defects (see FIG S1C for classification scheme). The percentage of cells with division defects is decreased by expression of *ltaS*^*F93L*^ (21.8%, n = 467) or *ltaS*^*L181S*^ (39.9%, n = 308) compared to expression of *ltaS*^*wt*^ (44.9%, n = 492). **G)** Purified sacculi from overnight RN4220 cultures were hydrolyzed with HCl and the released orthophosphate was quantified by an ammonium molybdate assay against a standard curve of KH_2_PO_4_. Δ*tarO* is a deletion mutant of the first gene in the WTA biosynthesis pathway and does not produce WTA. The average and standard deviation from 4 biological replicates are shown.

Access to a set of otherwise isogenic strains expressing different *ltaS* alleles allowed us to directly compare the impact of altering LTA length and abundance without confounding effects of other genetic differences. In the wild type background, we found that LTA was undetectable for *ltaS*^*Ins3*^; for the other two *ltaS* mutant alleles, LTA was shorter and the signal was much less intense than for cells expressing *ltaS*^*wt*^ (Fig. 4C). In the Δ*ugtP* background, LTA was nearly undetectable for *ltaS*^*Ins3*^ but appeared similar in length to *ltaS*^*wt*^. The other two mutants showed greatly reduced LTA signal intensity and appeared shorter in length than *ltaS*^*wt*^. Overall, these results confirmed that reducing LTA length or greatly reducing abundance in cells that produce long, abnormal LTA confers resistance to D-alanylation inhibitors. A question for future work is how D-alanylation protects cells from the lethality of long LTA polymers.

### Suppressors with low LTA retain the ability to produce large amounts of DAG

We noted that most of the *ltaS* mutant genes with suppressor mutations maintained the reading frame, yet the LTA levels in the mutants were typically low. LtaS produces DAG in addition to LTA, with one molecule of DAG produced during each round of elongation (Fig. 1B-C). DAG is generally thought to be a byproduct of LTA synthesis that serves mainly as a feedstock for glycolipid synthesis and for a recycling pathway that leads to more Ptd-Gro (53-55). To verify that the *ltaS* mutants retained the ability to produce appreciable amounts of DAG and to assess the correlation between DAG and LTA levels, we purified and reconstituted LtaS^wt^, LtaS^Ins3^, LtaS^F93L^, and LtaS^L181S^ in proteoliposomes and quantified DAG production (Fig. S6A). As reported previously, LtaS^wt^ made abundant LTA polymer in this system (56), but LTA was low to undetectable for the three mutants (Fig. S6B). However, all the mutants retained the capability to produce considerable amounts of DAG, ranging from ∼20% of LtaS^wt^ levels for LtaS^Ins3^ to levels comparable to LtaS^wt^ for LtaS^L181S^ (Fig. S6C). Because these are technically challenging assays, we only measured endpoints so relative DAG levels should be interpreted cautiously. Nonetheless, and with this caveat in mind, it appeared that the mutants produced more DAG than expected based on the LTA polymer detected, suggesting that DAG production may be partially uncoupled from LTA synthesis.

### Reducing LTA length and abundance partially corrects cell size and division defects and protects against peptidoglycan hydrolases

With our set of otherwise isogenic strains expressing *ltaS* variants, we next examined whether the *ltaS* suppressor mutations corrected other defects of Δ*ugtP.* We found that the rebuilt strain expressing *ltaS*^*wt*^ in the Δ*ugtP* background was highly susceptible to both Sle1 and lysostaphin, but strains expressing *ltaS* variants were not (Fig. 4D, Fig. S7). We also quantified cell morphology by microscopy and found that cells expressing *ltaS*^*F93L*^ or *ltaS*^*L181S*^ in the Δ*ugtP* background were 46% and 88% as large, respectively, as cells expressing *ltaS*^*wt*^ (Fig. 4E). These mutants had correspondingly fewer cell division defects (Fig. 4F). Although *ltaS*^*F93L*^ and *ltaS*^*L181S*^ produce LTA of similar length, *ltaS*^*F93L*^ corrected size and division defects to a greater extent compared to *ltaS*^*L181S*^. We attribute this observation to the difference in LTA abundance produced by these two strains. The abundance of LTA in *ltaS*^*F93L*^ more closely matched the abundance of LTA in the wild type strain, suggesting that the limited amount of LTA in *ltaS*^*L181S*^ may be insufficient to promote proper growth and division, as seen in *ltaS* depletion strains that grow abnormally large before lysing (5). Therefore, there may be both an ideal LTA length and abundance for optimal growth. The *ltaS*^*Ins3*^ mutant could not be quantified due to severe clumping and morphological aberrations. Nevertheless, the results for the other mutants allow us to conclude that reducing LTA length and abundance partially corrects the cell size and division defects of Δ*ugtP* mutant cells and also confers protection against PG hydrolases.

### Suppressors with low LTA have high WTA levels

Despite producing LTA at levels substantially below wild type levels, the *ltaS* mutants we selected were viable and relatively healthy. Moreover, two grew similarly to the wild type (Fig. 4B). Because LTA and WTA have partially overlapping roles, we hypothesized that there may be compensatory changes in WTA abundance in the *ltaS* mutants that contribute to their fitness. To address this question, we treated purified sacculi with strong acid to release orthophosphate from WTA and quantified it. We found that sacculi from strains expressing the mutant *ltaS* alleles contained two to three times as much phosphate as sacculi from the strain expressing *ltaS*^*wt*^, consistent with a large increase in WTA (Fig. 4G). PAGE analysis of WTA polymers liberated from purified sacculi with base showed modest increases in WTA length in the strains expressing *ltaS* mutant alleles (Fig. S8). We conclude that *S. aureus* adapts to a reduction in LTA by increasing WTA. Previous work has shown that WTA positively regulates cell wall crosslinking (26). Moreover, WTA is known to decrease susceptibility to PG hydrolases (23, 25). Therefore, we surmise that increased WTA levels and the concomitant changes in cell wall structure lead to the decreased susceptibility of our *ltaS* mutant strains to cell wall hydrolases.

## DISCUSSION

The principal goals of this investigation were to elucidate the roles of *ugtP* and LTA length in *S. aureus* cell growth and division and to identify mechanisms contributing to cell viability when LTA levels are low. First, by comparing multiple phenotypes of Δ*ltaA* and Δ*ugtP* mutants, we showed that cells producing long, abnormal LTA have cell growth and division defects and are also more susceptible to cell lysis than the wild type. Second, we showed that suppressor mutations in *ltaS* that reduced LTA length and abundance corrected these defects. Third, by comparing otherwise isogenic *ltaS* mutants that make either high or low levels of LTA, we showed that WTA levels correlated inversely with LTA levels. It is likely that the increased abundance of WTA partially compensated for low levels of LTA.

Previous work has shown that *S. aureus* Δ*ugtP* mutants, which make long LTA, grow larger than normal (32). In *B. subtilis, ugtP* is also important in cell growth and division, although its absence leads to decreased cell size in some reports (37, 38). *B. subtilis* UgtP is proposed to function as a nutrient sensor that, when present with sufficient UDP-glucose, slows Z-ring assembly so that cells grow to a larger size prior to division. Our results show that such a mechanism does not operate in *S. aureus.* First, Δ*ugtP* mutant cells spend a longer time growing before initiating septal synthesis and consequently are larger rather than smaller than the wild type. Second, Δ*ltaA* mutants are also larger than wild type even though they express *ugtP* and produce intracellular Glc_2_DAG. Therefore, dysregulated cell growth in LTA pathway mutants upstream of the LtaS polymerase is not due to the absence of the proteins or their products, but rather to the production of abnormally long LTA. Consistent with this conclusion, cell growth defects were reduced in a Δ*ugtP* mutant background by mutations in *ltaS* that reduced LTA length.

Previous work has shown that cell division defects result when *S. aureus* cells grow larger than normal (6, 42-45). Consistent with these findings, we observed a remarkable correlation between cell size and frequency of cell division defects for the mutants studied here. For example, Δ*ltaA* mutants were substantially smaller than Δ*ugtP* mutants, and we found that Δ*ltaA* mutants had 50% fewer cell division defects. Moreover, *ltaS* mutations that reduced cell size in a Δ*ugtP* background had fewer cell division defects, with the magnitudes of the reduction in size and the reduction in division defects tracking closely. We have therefore concluded that the cell division defects observed in Δ*ugtP* and Δ*ltaA* mutants are due to dysregulated coordination between cell growth and division.

We have shown that *S. aureus* WTA levels increase substantially when LTA levels are low. This finding is reminiscent of findings in *Streptococcus pneumoniae* showing that levels of WTA and LTA are inversely regulated (57). *S. pneumoniae* synthesizes both forms of teichoic acid through the same pathway, with the outcome distinguished only by a final ligation step where the polymer precursor is transferred to PG or to a glycolipid (58). In *S. pneumoniae*, LTA synthesis predominates during exponential growth, but there is a switch to predominantly WTA synthesis as cells approach stationary phase. *S. aureus* WTA and LTA are synthesized by different pathways with only the D-alanine tailoring modification as a shared feature. Nevertheless, there appears to be a mechanism to redistribute resources between the WTA and LTA pathways when one is lacking. Although LTA and WTA share some functions and act synergistically to maintain cell envelope integrity, their spatial localization is different. LTA is located between the membrane and the inner layer of PG and WTA extends from the inner layer of PG beyond the outer layer. Consistent with this difference in localization, we have found that producing long, abundant LTA tends to promote cell lysis whereas producing abundant WTA tends to limit cell lysis. Given this evidence that physiological roles of LTA and WTA in *S. aureus* are not identical, it would not be surprising to find that regulatory mechanisms exist to control the relative abundance of these polymers.

Expression of normal LTA is critical for *S. aureus* virulence. Previous work has shown that strains lacking *ltaA* or *ugtP* have attenuated pathogenicity (10, 11). While the physiological basis of this attenuation is not known, in light of our data on the susceptibility of Δ*ltaA* and Δ*ugtP* to PG hydrolases, it may involve an increased sensitivity to host lytic enzymes. We have shown here that the defects caused by expression of long, abnormal LTA also result in increased susceptibility of MRSA to beta-lactam antibiotics. We therefore anticipate that inhibitors of enzymes that act upstream of LtaS will have therapeutic potential, particularly if used in combination with an appropriate beta-lactam. Future work will focus on identifying such inhibitors and elucidating the mechanism for glycolipid-dependent control of LTA polymer length.

## MATERIALS AND METHODS

### General information

Chemicals and reagents were purchased from Sigma-Aldrich unless otherwise indicated. Oligonucleotides were purchased from Integrated DNA Technologies. Detergents were purchased from Anatrace. Restriction enzymes were purchased from New England Biolabs. *Staphylococcus aureus* strains were grown with shaking at 30°C unless otherwise indicated in tryptic soy broth (TSB, Beckton Dickinson Biosciences) or cation-adjusted Mueller Hinton Broth 2 (MHB2). *Escherichia coli* strains were grown at 37°C with shaking in LB broth, Miller (LB, Beckton Dickinson Biosciences). Growth on solid medium used the appropriate broth plus 1.5% w/v agar (Beckton Dickinson Biosciences). When required, antibiotics were used at the following concentrations: 50 μg/mL kanamycin, 50 μg/mL neomycin, 10 μg/mL erythromycin, 10 μg/mL chloramphenicol, 3 μg/mL tetracycline, 100 μg/mL carbenicillin. Anhydrotetracyline was used at 400 nM. *S. aureus* genomic DNA was isolated using a Wizard Genomic DNA Purification Kit (Promega). Genomic DNA from *S. aureus* RN4220 or isolated amsacrine-resistant mutants was used as a template in PCR reactions to amplify *S. aureus* genes.

### Strain construction

To construct strains with atc-inducible, integrated copies of genes via pTP63, plasmids were electroporated into *S. aureus* RN4220 containing the pTP44 plasmid. Transformants were selected on chloramphenicol at 30°C. Marked deletions, marked transposon insertions, and atc-inducible genes were transduced via ϕ85 bacteriophage as described previously (59).

### Analysis of LTA length by Western blot

Overnight cultures grown in TSB were diluted in fresh TSB and grown to an approximate OD_600_ of 0.8. 0.5 mL of each culture (normalized by OD_600_) were harvested by centrifugation and suspended in 50 μL of a solution containing 50 mM Tris pH 7.4, 150 mM NaCl, and 200 µg/mL lysostaphin (from *Staphylococcus staphylolyticus*, AMBI Products). The cells were incubated at 37°C for 10 minutes, diluted with one volume of 4X sodium dodecyl sulfate (SDS)-PAGE loading buffer, and boiled for 30 minutes. Samples were centrifuged for 10 minutes at 16,000 x g to pellet any insoluble material. The supernatant was diluted with one volume of water and treated with 0.5 μL of 20 mg/mL proteinase K (New England Biolabs) at 50°C for 2 hours. Samples were separated on 4-20% Mini-PROTEAN TGX acrylamide gels (Bio-rad) with a running buffer consisting of 5 g/L Tris base, 15 g/L glycine, and 1 g/L SDS and transferred to a PVDF membrane. Western blots were performed as described previously (56).

### Transmission electron microscopy

Overnight cultures grown in TSB were diluted in fresh TSB and grown to mid-log phase. Cells were treated with DMSO or 16 μg/mL amsacrine for 4 hours at 30°C, added to an equal volume of fixative solution (1.25% formaldehyde, 2.5% glutaraldehyde, 0.03% picric acid in 100 mM sodium cacodylate pH 7.4), and pelleted for fixation. Samples were prepared for TEM by the Harvard Medical School Electron Microscopy Facility and images were captured on a JEOL 1200EX instrument.

### Microscopy

Overnight cultures grown in TSB were diluted in fresh TSB and grown to an approximate OD_600_ of 0.5. To stain the cell membrane, FM 4-64 (ThermoFisher Scientific) was added at a final concentration of 5 μg/mL to 1 mL of culture for 5 minutes at room temperature. Cells were pelleted at 4,000*g* for 1 minute, washed with PBS pH 7.4 (50 mM sodium phosphate pH 7.4, 150 mM NaCl), and suspended in 100 μL of PBS pH 7.4. A 1 μL aliquot of cells was spotted on top of a 2% agarose gel pad mounted on a glass slide. A 1.5 mm cover slip was placed over the cells and sealed with wax before imaging.

The cells were imaged at 30°C as described previously (45). For each field of view, 3 images were taken: 1) phase-contrast, 2) brightfield, and 3) fluorescence. The phase-contrast and brightfield images were acquired at 100 ms camera exposure while the fluorescence image was acquired at 500 ms. The brightfield images were used for cell segmentation for quantitative image analyses. Fluorescence images were used to detect division defects and sort cells as depicted in (FIG S1C).

Image segmentation, cell volume quantification, and cellular phenotype classification were performed as described previously (45). Cell volumes were calculated from cells lacking visible septa. 600-1000 cells of each strain were quantified. A two-tailed Mann-Whitney U nonparametric test was used to calculate the p-value for the differences in distribution of cell sizes among strains. 300-600 cells of each strain were assessed for cell division phenotypes.

### Expression and purification of MBP-Sle1

*S. aureus* Sle1 (SAOUHSC_00427) was cloned into the NdeI and BamHI sites of pMAL-c5X (New England Biolabs) with primers oTD22 and oTD23 and transformed into *E. coli* NEB Express cells. An overnight culture grown at 30°C was used to inoculate fresh media and the culture was grown at 37°C to an approximate OD_600_ of 0.5. Cultures were cooled on ice and induced with 0.3 mM IPTG at 16°C overnight. Cells were pelleted at 3,600*g* for 15 minutes at 4°C, suspended in lysis buffer (20 mM Tris pH 7.2, 200 mM NaCl, 10% glycerol) plus 1 mM dithiothreitol, 1 mM phenylmethylsulfonyl fluoride, 100 μg/mL lysozyme, and 100 μg/mL DNase, and lysed with 3 passages through an EmulsiFlex-C3 cell disruptor (Avestin). All subsequent steps were performed at 4°C. Insoluble material was pelleted at 119,000*g* for 35 minutes and the supernatant was bound to amylose resin (New England Biolabs). The resin was washed with lysis buffer and eluted with lysis buffer plus 10 mM maltose. The elution was concentrated with a 30 kDa MWCO spin concentrator (EMD Millipore) and further purified on a Superdex 200 Increase 10/300 GL (GE Life Sciences) equilibrated in lysis buffer. Appropriate fractions were concentrated, flash frozen with liquid nitrogen, and stored at −80°C.

### Whole cell lytic assays

Overnight cultures grown in TSB were diluted in fresh TSB and grown to an approximate OD_600_ of 3. 1.5 mL of culture (normalized by OD_600_) were harvested by centrifugation, washed with PBS pH 7.2, and suspended in 1.6 mL of PBS pH 7.2. 75 μL of cell suspension were added to 75 μL of 25 nM lysostaphin, 250-500 nM purified Sle1, or PBS pH 7.2. Samples were incubated at 25°C with shaking in a SpectaMax Plus 384 microplate reader and OD_600_ was monitored over time.

### Minimum inhibitory concentration (MIC) determination

Overnight cultures grown in TSB were diluted in fresh MHB2 (without antibiotics) and grown to an approximate OD_600_ of 0.8. Cultures were diluted with media to an OD_600_ of 0.001 and 146-147 μL were added to each well of a 96-well plate. 3-4 μL of a compound dilution series were added, and cultures were grown with shaking at 37°C for 18 hours. Each condition was tested in technical triplicates and the MIC was determined as the lowest concentration that prevented growth.

### Spot dilutions

Overnight cultures of each strain grown in TSB were diluted in fresh TSB and grown to an approximate OD_600_ of 0.8. Cultures were normalized to an OD_600_ of 0.1, a 10-fold dilution series from 10^−1^ to 10^−6^ was created, and dilutions were spotted on TSB agar containing any desired compounds. Strains growth with anhydrotetracycline were washed once with TSB before diluting. Plates were incubated at 30°C and imaged the next day with a Nikon D3400 DSLR camera fitted with an AF Micro-Nikkor 60 mm f/2.8D lens.

### Growth curves

Overnight cultures of each strain grown in TSB were diluted in fresh TSB and grown to an approximate OD_600_ of 0.8. Cultures were diluted to an OD_600_ of 0.03 and amsacrine or DBI-1 (in DMSO) were added at a final concentration of 10 μg/mL. DMSO was added to untreated control cultures at a final concentration of 2%. Cultures were grown at 30°C with shaking in a SpectaMax Plus 384 microplate reader (Molecular Devices) and OD_600_ was monitored over time.

### Raising amsacrine resistant mutants

Amsacrine resistant mutants were raised strains SEJ1 *ugtP::tn*, HG003 *ugtP::tn*, RN4220 Δ*ugtP*, and SEJ1 Δ*ugtP*::*kan*. For mutants in the SEJ1 *ugtP::tn* background, 50 µL of undiluted overnight cell culture was plated on TSB agar plus 6 µg/mL amsacrine at 30°C for two days. For mutants in backgrounds SEJ1 Δ*ugtP*::*kan*, HG003 *ugtP::tn*, RN4220 Δ*ugtP*, overnight cultures were diluted in TSB and grown at 30°C to an OD_600_ of 1.0. 1 mL of this culture was harvested, suspended in 100 µL fresh TSB, and plated on TSB agar plus 10 µg/mL amsacrine at 30°C for two days. Multiple independent cultures were used to increase the diversity of mutants. Whole-genome sequencing of select mutants was performed with an Illumina MiSeq as described previously (45).

### Construction of strains with anhydrotetracycline-inducible *ltaS* alleles

Wild type, Ins3, F93L, and L181S LtaS variants were cloned from the genomic DNA of RN4220 or their respective suppressor mutants into pTP63 with primers iLtaS_1 and iLtaS_2 and electroporated into RN4220 bearing pTP44 for integration (44). Each resulting strain was transduced with ϕ85 lysate from a strain with an erythromycin-marked *ltaS* deletion. These strains were then optionally transduced with ϕ85 lysate from a strain with a kanamycin-marked *ugtP* or *ltaA* deletion.

### Phosphate quantification from purified sacculi

Sacculi containing covalently linked WTA were isolated in a manner similar as described previously (60). 2 mL of an overnight culture grown in TSB (normalized by OD_600_) were harvested, washed with buffer 1 (50 mM MES pH 6.5), and suspended in buffer 2 (50 mM MES pH 6.5, 4% SDS). Cells were boiled for 1 hour and pellets harvested at 16,000*g* for 10 minutes. The supernatant was discarded, and the pellets were washed twice with buffer 2, once with buffer 3 (50 mM MES pH 6.5, 342 mM NaCl), and twice with buffer 1. Pellets were treated with 50 μg/mL DNase and 50 μg/mL RNase in buffer 1 at 37°C for 1 hour. Pellets were harvested, washed with buffer 1, and suspended in a solution containing 20 mM Tris pH 8, 0.5% SDS. Samples were treated with 20 μg/mL proteinase K at 50°C for 2 hours with light shaking. After harvesting pellets by centrifugation, pellets were washed once with buffer 3 and then 3 times with water.

Purified sacculi were suspended in 1M HCl. A dilution series of K_2_HPO_4_ in 1M HCl was also prepared. Samples were treated at 80°C for 16 hours and cooled to room temperature. Any insoluble material remaining was pelleted by centrifugation and the supernatant was retained. An ammonium molybdate reagent was prepared by mixing, in order, equal volumes of 2M H_2_SO_4_, 2.5% (w/v) ammonium molybdate, and 10% (w/v) ascorbic acid. One volume of ammonium molybdate reagent was added to each sample and samples were incubated at 37°C for 1 hour. Orthophosphate was quantified by absorbance at 820nm with the K_2_HPO_4_ standard curve.

### Polyacrylamide gel electrophoresis of WTA polymers

Sacculi containing covalently linked WTA were isolated as described above but treated with 100 mM NaOH at room temperature for 16 hours. 3 volumes of loading buffer (50% glycerol, 100 mM Tris-tricine, 0.02% bromophenol blue) were added to each sample.

High-resolution 20×20cm polyacrylamide gels were prepared as described previously (60), but with a stacking gel consisting of 3% acrylamide (3% T, 3.3% C where T is total acrylamide and C is the percentage of T consisting of bisacrylamide), 900 mM Tris pH 8.5, 0.1% ammonium persulfate, and 0.01% tetramethylethylenediamine. Gels were run at 4°C in a Protean II xi Cell electrophoresis system (Bio-rad) at 40 mA/gel with a running buffer consisting of 100 mM Tris-tricine pH 8.2 until the bromophenol blue loading dye was near the bottom of the gel. Gels were washed with water, stained with 1 mg/mL aqueous alcian blue for 30 minutes, destained with water and 40% ethanol/5% acetic acid, and rehydrated with water. Silver stain was performed with the Silver Stain Plus kit (Bio-rad) without the fixation step. Images were taken with a Nikon D3400 DSLR camera fitted with an AF Micro-Nikkor 60 mm f/2.8D lens and converted to an 8-bit image using ImageJ.

### Purification of LtaS mutants and proteoliposome analysis of DAG production

Mutant LtaS constructs were cloned from genomic DNA isolated from the original suppressor mutants and assembled into pET28b with primers LtaS_F and LtaS_R as previously described (56). LtaS constructs were expressed, purified, and reconstituted into proteoliposomes as previously described (56). Proteoliposomes were added to 9 volumes of a solution containing 20 mM succinate 6.0, 50 mM NaCl, 5% DMSO. Reactions were incubated either in the presence or absence of 1 mM MnCl_2_. LTA was detected by Western blot as previously described (56). To measure DAG production, reactions proceeded for 4 hours at 30°C, and then extracted and assayed according to the instructions provided by the Cell BioLabs DAG assay kit. Reactions were performed in duplicate and plotted using GraphPad Prism. Absolute activity was calculated by subtracting activity values calculated from reactions that did not contain MnCl_2_ from the values from reactions that contained MnCl_2_. Activity was compared between mutants by setting reactions with proteoliposomes containing wild type LtaS to 100% activity.

### Data availability

Whole-genome sequencing data (accession number PRJNA612838) can be found in the NCBI BioProject database.

## SUPPLEMENTAL MATERIAL

**Supplemental File 1**, PDF file, 1.8 MB.

## ACKNOWLEDGMENT

Some strains were generated with the help of Lincoln Pasquina and Wonsik Lee and we thank them for their generous assistance. This project was funded by the National Institutes of Health (P01-AI083214 to S.W. and R01-AI139011 to S.W. and R.L.) and the National Science Foundation (DGE1144152 to A.R.H. and to T.D.).

## REFERENCES

1. Rajagopal M, Walker S. 2017. Envelope Structures of Gram-Positive Bacteria. Curr Top Microbiol Immunol 404:1–44.

2. Santa Maria JP, Jr., Sadaka A, Moussa SH, Brown S, Zhang YJ, Rubin EJ, Gilmore MS, Walker S. 2014. Compound-gene interaction mapping reveals distinct roles for Staphylococcus aureus teichoic acids. Proc Natl Acad Sci U S A 111:12510–5.

3. Oku Y, Kurokawa K, Matsuo M, Yamada S, Lee B-L, Sekimizu K. 2009. Pleiotropic Roles of Polyglycerolphosphate Synthase of Lipoteichoic Acid in Growth of Staphylococcus aureus Cells. Journal of Bacteriology 191:141–151.

4. Schirner K, Marles-Wright J, Lewis RJ, Errington J. 2009. Distinct and essential morphogenic functions for wall- and lipo-teichoic acids in Bacillus subtilis. The EMBO Journal 28:830–842.

5. Gründling A, Schneewind O. 2007. Synthesis of glycerol phosphate lipoteichoic acid in Staphylococcus aureus. Proc Natl Acad Sci U S A 104:8478–83.

6. Campbell J, Singh AK, Santa Maria JP, Jr., Kim Y, Brown S, Swoboda JG, Mylonakis E, Wilkinson BJ, Walker S. 2010. Synthetic Lethal Compound Combinations Reveal a Fundamental Connection between Wall Teichoic Acid and Peptidoglycan Biosyntheses in Staphylococcus aureus. ACS Chem Biol 6:106–116.

7. Weidenmaier C, Kokai-Kun JF, Kristian SA, Chanturiya T, Kalbacher H, Gross M, Nicholson G, Neumeister B, Mond JJ, Peschel A. 2004. Role of teichoic acids in Staphylococcus aureus nasal colonization, a major risk factor in nosocomial infections. Nat Med 10:243–5.

8. Weidenmaier C, Peschel A, Kempf VA, Lucindo N, Yeaman MR, Bayer AS. 2005. DltABCD- and MprF-mediated cell envelope modifications of Staphylococcus aureus confer resistance to platelet microbicidal proteins and contribute to virulence in a rabbit endocarditis model. Infect Immun 73:8033–8.

9. Wanner S, Schade J, Keinhörster D, Weller N, George SE, Kull L, Bauer J, Grau T, Winstel V, Stoy H, Kretschmer D, Kolata J, Wolz C, Bröker BM, Weidenmaier C. 2017. Wall teichoic acids mediate increased virulence in Staphylococcus aureus. Nat Microbiol 2:16257.

10. Gründling A, Schneewind O. 2007. Genes required for glycolipid synthesis and lipoteichoic acid anchoring in Staphylococcus aureus. J Bacteriol 189:2521–30.

11. Sheen TR, Ebrahimi CM, Hiemstra IH, Barlow SB, Peschel A, Doran KS. 2010. Penetration of the blood-brain barrier by Staphylococcus aureus: contribution of membrane anchored lipoteichoic acid. Journal of molecular medicine (Berlin, Germany) 88:633–639.

12. Collins LV, Kristian SA, Weidenmaier C, Faigle M, van Kessel KPM, van Strijp JAG, Götz F, Neumeister B, Peschel A. 2002. Staphylococcus aureus strains lacking D-alanine modifications of teichoic acids are highly susceptible to human neutrophil killing and are virulence attenuated in mice. J Infect Dis 186:214–219.

13. Simanski M, Gläser R, Köten B, Meyer-Hoffert U, Wanner S, Weidenmaier C, Peschel A, Harder J. 2013. Staphylococcus aureus subverts cutaneous defense by D-alanylation of teichoic acids. Exp Dermatol 22:294–6.

14. Bunk S, Sigel S, Metzdorf D, Sharif O, Triantafilou K, Triantafilou M, Hartung T, Knapp S, von Aulock S. 2010. Internalization and coreceptor expression are critical for TLR2-mediated recognition of lipoteichoic acid in human peripheral blood. J Immunol 185:3708–17.

15. Neuhaus FC, Baddiley J. 2003. A continuum of anionic charge: structures and functions of D-alanyl-teichoic acids in gram-positive bacteria. Microbiol Mol Biol Rev 67:686–723.

16. Corrigan RM, Abbott JC, Burhenne H, Kaever V, Gründling A. 2011. c-di-AMP is a new second messenger in Staphylococcus aureus with a role in controlling cell size and envelope stress. PLoS Pathog 7:e1002217.

17. Wang H, Gill CJ, Lee SH, Mann P, Zuck P, Meredith TC, Murgolo N, She X, Kales S, Liang L, Liu J, Wu J, Santa Maria J, Su J, Pan J, Hailey J, McGuinness D, Tan CM, Flattery A, Walker S, Black T, Roemer T. 2013. Discovery of Wall Teichoic Acid Inhibitors as Potential Anti-MRSA beta-Lactam Combination Agents. Chem Biol 20:272–84.

18. Schuster CF, Wiedemann DM, Kirsebom FCM, Santiago M, Walker S, Gründling A. 2019. High-throughput transposon sequencing highlights the cell wall as an important barrier for osmotic stress in methicillin resistant Staphylococcus aureus and underlines a tailored response to different osmotic stressors. Mol Microbiol doi: 10.1111/mmi.14433.

19. Peschel A, Otto M, Jack RW, Kalbacher H, Jung G, Gale RT, Götz F. 1999. Inactivation of the dlt operon in Staphylococcus aureus confers sensitivity to defensins, protegrins, and other antimicrobial peptides. J Biol Chem 274:8405–8410.

20. Kohler T, Weidenmaier C, Peschel A. 2009. Wall Teichoic Acid Protects Staphylococcus aureus against Antimicrobial Fatty Acids from Human Skin. Journal of Bacteriology 191:4482–4484.

21. Brown S, Xia G, Luhachack LG, Campbell J, Meredith TC, Chen C, Winstel V, Gekeler C, Irazoqui JE, Peschel A, Walker S. 2012. Methicillin resistance in Staphylococcus aureus requires glycosylated wall teichoic acids. Proc Natl Acad Sci U S A 109:18909–14.

22. Mishra NN, Bayer AS, Weidenmaier C, Grau T, Wanner S, Stefani S, Cafiso V, Bertuccio T, Yeaman MR, Nast CC, Yang SJ. 2014. Phenotypic and genotypic characterization of daptomycin-resistant methicillin-resistant Staphylococcus aureus strains: relative roles of mprF and dlt operons. PLoS One 9:e107426.

23. Frankel MB, Schneewind O. 2012. Determinants of murein hydrolase targeting to cross-wall of Staphylococcus aureus peptidoglycan. J Biol Chem 287:10460–71.

24. Fedtke I, Mader D, Kohler T, Moll H, Nicholson G, Biswas R, Henseler K, Götz F, Zähringer U, Peschel A. 2007. A Staphylococcus aureus ypfP mutant with strongly reduced lipoteichoic acid (LTA) content: LTA governs bacterial surface properties and autolysin activity. Molecular Microbiology 65:1078–1091.

25. Sieradzki K, Tomasz A. 2003. Alterations of cell wall structure and metabolism accompany reduced susceptibility to vancomycin in an isogenic series of clinical isolates of Staphylococcus aureus. J Bacteriol 185:7103–10.

26. Atilano ML, Pereira PM, Yates J, Reed P, Veiga H, Pinho MG, Filipe SR. 2010. Teichoic acids are temporal and spatial regulators of peptidoglycan cross-linking in Staphylococcus aureus. Proc Natl Acad Sci U S A 107:18991–6.

27. Tiwari KB, Gatto C, Walker S, Wilkinson BJ. 2018. Exposure of Staphylococcus aureus to Targocil Blocks Translocation of the Major Autolysin Atl across the Membrane, Resulting in a Significant Decrease in Autolysis. Antimicrob Agents Chemother 62.

28. D’Elia MA, Pereira MP, Chung YS, Zhao W, Chau A, Kenney TJ, Sulavik MC, Black TA, Brown ED. 2006. Lesions in teichoic acid biosynthesis in Staphylococcus aureus lead to a lethal gain of function in the otherwise dispensable pathway. J Bacteriol 188:4183–9.

29. Bæk KT, Bowman L, Millership C, Dupont Søgaard M, Kaever V, Siljamäki P, Savijoki K, Varmanen P, Nyman TA, Gründling A, Frees D. 2016. The Cell Wall Polymer Lipoteichoic Acid Becomes Nonessential in Staphylococcus aureus Cells Lacking the ClpX Chaperone. MBio 7.

30. Karinou E, Schuster CF, Pazos M, Vollmer W, Gründling A. 2019. Inactivation of the Monofunctional Peptidoglycan Glycosyltransferase SgtB Allows Staphylococcus aureus To Survive in the Absence of Lipoteichoic Acid. J Bacteriol 201.

31. Jorasch P, Warnecke DC, Lindner B, Zähringer U, Heinz E. 2000. Novel processive and nonprocessive glycosyltransferases from Staphylococcus aureus and Arabidopsis thaliana synthesize glycoglycerolipids, glycophospholipids, glycosphingolipids and glycosylsterols. Eur J Biochem 267:3770–3783.

32. Kiriukhin MY, Debabov DV, Shinabarger DL, Neuhaus FC. 2001. Biosynthesis of the glycolipid anchor in lipoteichoic acid of Staphylococcus aureus RN4220: role of YpfP, the diglucosyldiacylglycerol synthase. J Bacteriol 183:3506–14.

33. Lu D, Wörmann ME, Zhang X, Schneewind O, Gründling A, Freemont PS. 2009. Structure-based mechanism of lipoteichoic acid synthesis by Staphylococcus aureus LtaS. Proc Natl Acad Sci U S A 106:1584–1589.

34. Duckworth M, Archibald AR, Baddiley J. 1975. Lipoteichoic acid and lipoteichoic acid carrier in Staphylococcus aureus H. FEBS Lett 53:176–179.

35. Taron DJ, Childs III WC, Neuhaus FC. 1983. Biosynthesis of D-Alanyl-Lipoteichoic Acid: Role of Diglyceride Kinase in the Synthesis of Phosphatidylglycerol for Chain Elongation. J Bacteriol 154:1110–1116.

36. Jerga A, Lu YJ, Schujman GE, de Mendoza D, Rock CO. 2007. Identification of a soluble diacylglycerol kinase required for lipoteichoic acid production in Bacillus subtilis. J Biol Chem 282:21738–45.

37. Price KD, Roels S, Losick R. 1997. A Bacillus subtilis gene encoding a protein similar to nucleotide sugar transferases influences cell shape and viability. J Bacteriol 179:4959–4961.

38. Weart RB, Lee AH, Chien AC, Haeusser DP, Hill NS, Levin PA. 2007. A metabolic sensor governing cell size in bacteria. Cell 130:335–47.

39. Lazarevic V, Soldo B, Medico N, Pooley H, Bron S, Karamata D. 2005. Bacillus subtilis alpha-phosphoglucomutase is required for normal cell morphology and biofilm formation. Appl Environ Microbiol 71:39–45.

40. Matsuoka S, Chiba M, Tanimura Y, Hashimoto M, Hara H, Matsumoto K. 2011. Abnormal morphology of Bacillus subtilis ugtP mutant cells lacking glucolipids. Genes Genet Syst 86:295–304.

41. Chien AC, Zareh SK, Wang YM, Levin PA. 2012. Changes in the oligomerization potential of the division inhibitor UgtP co-ordinate Bacillus subtilis cell size with nutrient availability. Mol Microbiol 86:594–610.

42. Pinho MG, Errington J. 2003. Dispersed mode of Staphylococcus aureus cell wall synthesis in the absence of the division machinery. Mol Microbiol 50:871–81.

43. Jorge AM, Hoiczyk E, Gomes JP, Pinho MG. 2011. EzrA contributes to the regulation of cell size in Staphylococcus aureus. PLoS One 6:e27542.

44. Pang T, Wang X, Lim HC, Bernhardt TG, Rudner DZ. 2017. The nucleoid occlusion factor Noc controls DNA replication initiation in Staphylococcus aureus. PLoS Genet 13:e1006908.

45. Do T, Schaefer K, Santiago AG, Coe KA, Fernandes PB, Kahne D, Pinho MG, Walker S. 2020. Staphylococcus aureus cell growth and division are regulated by an amidase that trims peptides from uncrosslinked peptidoglycan. Nat Microbiol doi: 10.1038/s41564-019-0632-1.

46. Kajimura J, Fujiwara T, Yamada S, Suzawa Y, Nishida T, Oyamada Y, Hayashi I, Yamagishi J, Komatsuzawa H, Sugai M. 2005. Identification and molecular characterization of an N-acetylmuramyl-L-alanine amidase Sle1 involved in cell separation of Staphylococcus aureus. Mol Microbiol 58:1087–101.

47. Schindler CA, Schuhardt VT. 1964. Lysostaphin: a New Bacteriolytic Agent for the Staphlyococcus. Proc Natl Acad Sci U S A 51:414–421.

48. Robinson JM, Hardman JK, Sloan GL. 1979. Relationship Between Lysostaphin Endopeptidase Production and Cell Wall Composition in Staphylococcus staphylolyticus. J Bacteriol 137:1158–1164.

49. Pasquina L, Santa Maria JP, Jr., Wood BM, Moussa SH, Matano LM, Santiago M, Martin SE, Lee W, Meredith TC, Walker S. 2016. A synthetic lethal approach for compound and target identification in Staphylococcus aureus. Nat Chem Biol 12:40–5.

50. Peschel A, Vuong C, Otto M, Götz F. 2000. The D-Alanine Residues of Staphylococcus aureus Teichoic Acids Alter the Susceptibility to Vancomycin and the Activity of Autolytic Enzymes. Antimicrob Agents Chemother 44:2845–2847.

51. Matano LM, Morris HG, Wood BM, Meredith TC, Walker S. 2016. Accelerating the discovery of antibacterial compounds using pathway-directed whole cell screening. Bioorg Med Chem 24:6307–6314.

52. Fahmi T, Port GC, Cho KH. 2017. c-di-AMP: An Essential Molecule in the Signaling Pathways that Regulate the Viability and Virulence of Gram-Positive Bacteria. Genes (Basel) 8.

53. Matsuoka S, Hashimoto M, Kamiya Y, Miyazawa T, Ishikawa K, Hara H, Matsumoto K. 2011. The Bacillus subtilis essential gene dgkB is dispensable in mutants with defective lipoteichoic acid synthesis. Genes Genet Syst 86:365–376.

54. Parsons JB, Rock CO. 2013. Bacterial lipids: metabolism and membrane homeostasis. Prog Lipid Res 52:249–76.

55. Kuhn S, Slavetinsky CJ, Peschel A. 2015. Synthesis and function of phospholipids in Staphylococcus aureus. Int J Med Microbiol 305:196–202.

56. Vickery CR, Wood BM, Morris HG, Losick R, Walker S. 2018. Reconstitution of Staphylococcus aureus lipoteichoic Acid synthase activity identifies congo red as a selective inhibitor. J Am Chem Soc 140:876–879.

57. Flores-Kim J, Dobihal GS, Fenton A, Rudner DZ, Bernhardt TG. 2019. A switch in surface polymer biogenesis triggers growth-phase-dependent and antibiotic-induced bacteriolysis. Elife 8.

58. Heß N, Waldow F, Kohler TP, Rohde M, Kreikemeyer B, Gómez-Mejia A, Hain T, Schwudke D, Vollmer W, Hammerschmidt S, Gisch N. 2017. Lipoteichoic acid deficiency permits normal growth but impairs virulence of Streptococcus pneumoniae. Nat Commun 8:2093.

59. Lee W, Do T, Zhang G, Kahne D, Meredith TC, Walker S. 2018. Antibiotic combinations that enable one-step, targeted mutagenesis of chromosomal genes. ACS Infect Dis doi: 10.1021/acsinfecdis.8b00017.

60. Meredith TC, Swoboda JG, Walker S. 2008. Late-stage polyribitol phosphate wall teichoic acid biosynthesis in Staphylococcus aureus. J Bacteriol 190:3046–56.

